# The cellular Activating Protein 1 cFos regulates Influenza A virus replication

**DOI:** 10.1101/2025.08.12.669837

**Authors:** Antoine Gerodez, François E. Dufrasne, Olivier Denis, Mieke Steensels, Bénédicte Lambrecht, Lionel Tafforeau, Caroline Demeret, Cyril Barbezange

## Abstract

Previous research has demonstrated that Influenza A virus (IAV) infection activates AP-1 transcription factors as part of the antiviral response. In this study, we identified cFos as the most upregulated AP-1 transcription factor during IAV infection in A549 human lung cells. Surprisingly, the knockdown of cFos resulted in impaired IAV replication. Fluorescence microscopy and functional analyses indicated that cFos is implicated in IAV infection through its nuclear function, rather than its cytoplasmic role as an activator of lipid synthesis. The investigation into the role of cFos in IAV infection revealed increased apoptosis and elevated interferon-β mRNA levels in cFos-knockdown A549 cells during IAV infection. This suggests that cFos may enhance cell survival and reduce interferon-β expression during infection, thereby facilitating IAV proliferation. Furthermore, the levels of viral NA mRNA and the expression of late viral proteins NA and M2 decreased upon cFos knockdown. Overall, this study identifies cFos as a proviral factor for IAV, through the modulation of innate immunity and apoptosis during infection, and potentially by supporting the viral transcription.

**Author Summary:** Influenza A viruses pose a significant threat to global public health. Understanding the interaction between the virus and host is essential for the development of antiviral therapies. We identified cFos, a cellular Activating Protein-1, as a pro-viral factor for influenza A viruses. cFos was found to enhance cell survival and regulate the expression of certain cytokines during viral infection. Our findings also suggested a potential role for cFos in the viral transcription, especially of the viral proteins expressed late in the viral cycle.

## Introduction

Activating Protein-1 (AP-1) transcription factors consist of homo- and hetero-dimers composed of basic region-leucine zipper (bZIP) proteins that belong to Jun (cJun, JunB, JunD), Fos (cFos, FosB, Fra-1, Fra-2), Maf (c-Maf, MafB, MafA, MafG/F/K and Nrl) and ATF (ATF2, LRF1/ATF3, B-ATF, JDP1, JDP2) families. These transcription factors regulate many cellular functions, including cell proliferation, differentiation, inflammation, and apoptosis (reviewed in refs. [1,2]). cFos and cJun are commonly associated in a complex that binds in a sequence-specific manner to the promoter and enhancer regions of target genes [3].

Although Jun and Fos families have been considered to regulate cell proliferation positively (reviewed in ref. [4]), the role of cFos in cell proliferation remains debatable. While mouse fibroblasts deficient in both cFos and FosB had a reduced proliferative activity, the inhibition of cFos alone induced no effect on cell [5]. In addition, double-knockout mice lacking both cFos and FosB, but not the single knockout mice, had smaller body sizes than their wild-type counterparts [6]. On the other hand, the knockdown of cFos resulted in inhibited proliferation of a human osteosarcoma cell line, and the stable overexpression of cFos led to increased proliferation of immortalized human hepatocytes under low serum conditions [7,8]. cFos also regulates apoptosis. Although cFos was shown to promote apoptosis in different cell types [9–11], it seemed to repress apoptosis in a human osteosarcoma cell line and to decrease neuronal cell death in the hippocampus during kainic-acid-induced seizure [7,12]. Therefore, the regulation of the cell fate by cFos and the AP-1, in general, is complex and depend on the cell type, the type and duration of the stimulus, and the involvement of other transcription factors. In addition, AP-1 has been described as an activator of inflammation (reviewed in ref. [2]), although some evidence also suggests a specific anti-inflammatory role of cFos. A significant increase in the pro-inflammatory cytokines such as TNF-α, IL-6 and IL-12 p40 were observed in mouse macrophages and mice deficient in cFos [13], while the expression of cFos was observed to inhibit IL-12 p40 promoter activity in mouse macrophages [14]. Finally, independently of its transcriptional factor activity in the nucleus, cFos possesses a cytoplasmic function as an activator of lipid synthesis at the endoplasmic reticulum level (reviewed in ref [15]).

Several viruses were shown to hijack AP-1 proteins to support their replication. AP-1 binding to the intragenic regulatory region of the *pol* gene of the human immunodeficiency virus type 1 (HIV-1) seemed to help recruiting the cellular DNA-dependent RNA-polymerase II to the viral promoter, supporting viral transcription [16]. cFos was found to bind to multiple gene promoters of Kaposi’s Sarcoma-associated herpesvirus and to enhance viral lytic transcription [17]. siRNA-based experiment identified a role of cFos in hepatitis C virus replication and propagation [18]. cFos was also shown to promote virus replication of alpha- and gamma-coronavirus by delaying and reducing apoptosis [19].

Influenza A viruses (IAVs) are major respiratory pathogens responsible for human seasonal epidemics and pandemics, posing a persistent threat to global health. IAVs are enveloped viruses and belong to the *Orthomyxoviridae* family [20]. Their genome consists of 8 negative-sense single-strand RNA segments. Each viral RNA (vRNA) segment is encapsidated by nucleoproteins (NP) and attached to the heterotrimeric polymerase complex, composed of the PB1, PB2, and PA subunits, thus forming the viral ribonucleoprotein (vRNP). To start an infection, IAV enters the cell through an endocytic pathway. Inside the cell, vRNPs are released and transported to the nucleus where viral transcription and replication occur. Transcription and replication are carried out by the viral polymerase complex. Primary transcription generates viral mRNAs, which are exported to the cytoplasm for translation by host ribosomes. PB2, PB1, PA, NP, and NS1 are expressed early while HA, NA, M1, and NS2 are expressed later during the viral cycle [21]. Viral proteins PB2, PB1, PA, NP, M1, and NS2 are transported back to the nucleus, and genome replication then occurs. Newly synthesized vRNAs are assembled with PB2, PB1, PA, and NP, resulting in progeny vRNPs that are subsequently exported to the cytoplasm. These vRNPs are further incorporated into progeny particles containing HA, NA, M2 and M1 inserted in or present at the cell membrane. Finally, progeny virions are released from the cell by budding for subsequent infection [20,22].

AP-1 transcription factors might play a role in IAV polymerase activity regulation, as the inhibition of cJun was shown to impair IAV H5N1 replication in human lung cells [23]. However, AP-1 also appears to take part in the innate antiviral response following IAV infection. IAV-induced AP-1 activation was shown to activate the expression of interferon-β and promote NLRP3 inflammasome activation [24,25]. Furthermore, the viral protein NS1 antagonized IAV-induced AP-1 activation [26]. Thus, the role of the AP-1 transcription factors in IAV infection remains unclear. In this study, we observed that the infection of human cells with human IAVs resulted in the upregulation of cFos and cJun subunits. The role of cFos and cJun was then investigated using specifically depleted cells, showing that knockdown of cFos —but not cJun— significantly impaired viral replication efficiency. Further characterization revealed different potential mechanisms by which cFos may support IAV replication.

## Results

### IAV multiplication upregulates the AP-1 transcription factors and is impaired in cFos-knockdown cells

Following single-cycle IAV infection in A549 cells, the expression of AP-1 transcription factors cFos, FosB, cJun, and JunD was upregulated at 6 and 9 hpi, as measured by RT-qPCR. The mRNA level of cFos and FosB was > 50-fold higher than in mock-infected cells. For cJun and JunB, the mRNA level at 9 hpi was between 5-10 times higher in infected cells. No difference in the mRNA expression of Fra1, Fra2, JunD, and ATF2 was observed at any time point (Fig 1A). The role of cFos and cJun on viral replication was further investigated using small interfering RNA (siRNA)-mediated silencing. Upon individual siRNA knockdown of cFos and cJun, A549 cell viability remained above 90% compared to the non-target (NT)-siRNA treated cells, with knockdown efficiency of protein expression estimated at 92 and 61% for cFos and cJun, respectively (S1 Fig A-B). cFos knockdown significantly impaired the replication of human seasonal pH1N1, H3N2, and WSN IAV strains. In contrast, IAV replication was not affected by the individual knockdown of cJun (Fig 1B). Knockdown of the well-described proviral factor RAB11 [27] was used as a control (S1 Fig C).

**Fig 1.**
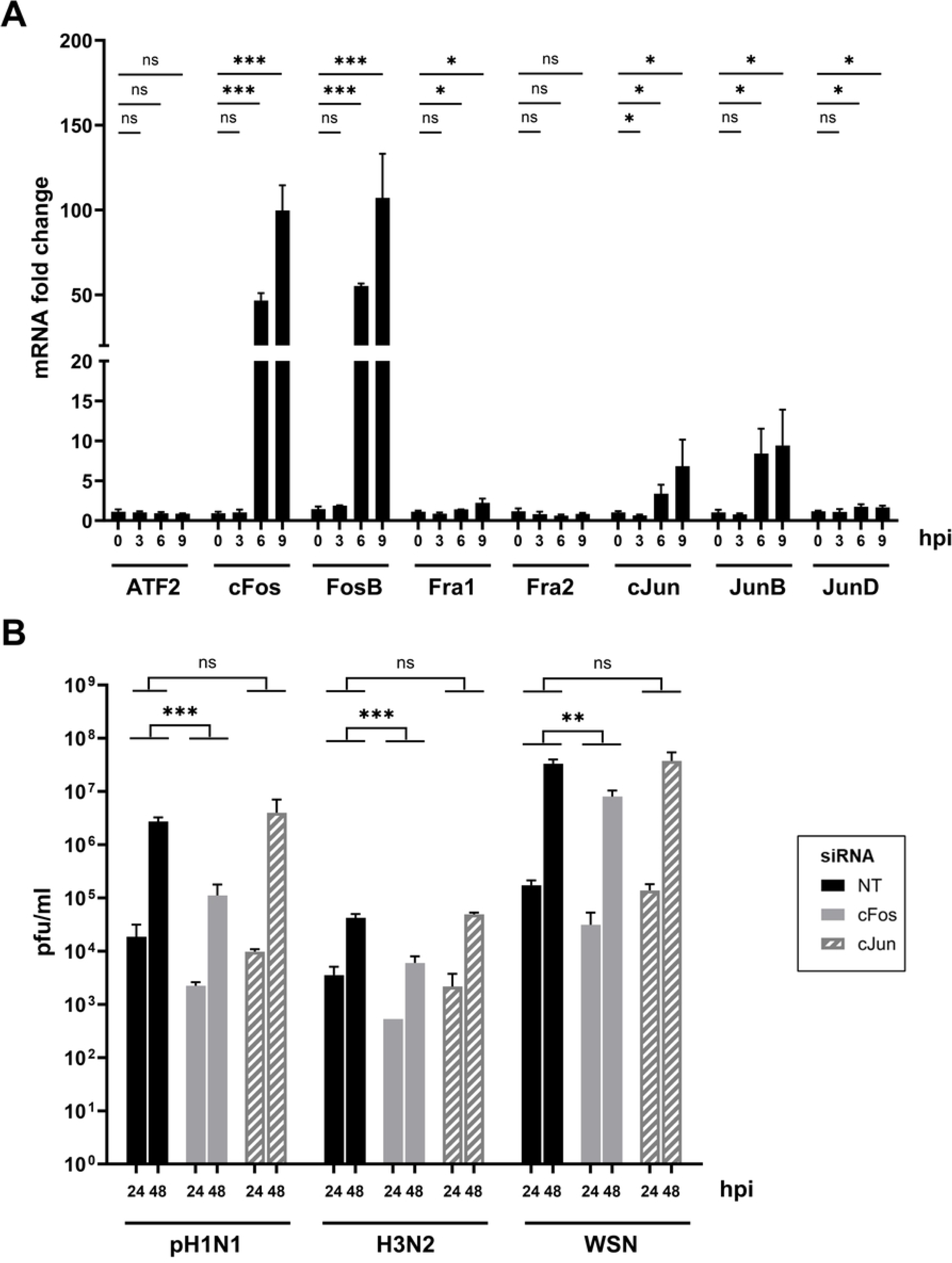
cFos mRNA expression is upregulated during IAV infection, and its knockdown reduced IAV replication. **(A)** A549 cells were infected with WSN at moi of 3 pfu/cell. Total RNAs were extracted at the indicated time point post-infection and the mRNA expression levels of AP-1 were determined by RT-qPCR. mRNA fold changes were calculated using the 2^-ΔΔCt^ method compared to the mock cells at each time point. Results are expressed at the mean ± SD determined in three independent experiments. The significance of the difference to the 0 hpi timepoint was tested with unpaired t-tests using GraphPad Prism software for each cellular factor (ns: not significant, *p< 0.05, **p< 0.01, ***p< 0.001). **(B)** A549 cells were transfected with 25 nM of the non-target (NT) or the indicated siRNAs. At 48 hpt, cells were infected with the following viruses at the indicated moi in pfu/cell: A/Bretagne/7608/2009(H1N1pdm09) (pH1N1, moi of 10^−3^); A/Centre/1003/2012(H3N2) (H3N2, moi of 10^−2^); A/WSN/33(H1N1) (WSN, moi of 10^−4^). At 0, 24 and 48 hpi, viral titers were determined by plaque-forming assay. Results are expressed as the mean ± SD pfu/ml of three independent experiments. The area under the curve (AUC) (not shown) was determined for each virus and condition, taking the pfu/ml at timing 0 hpi as the baseline. The significance was tested on AUCs with unpaired t tests using GraphPad Prism software (ns: not significant, *p< 0.05, **p< 0.01, ***p< 0.001).

### The nuclear function of cFos, but not its cytoplasmic function, regulates IAV multiplication

We were further interested in deciphering which function of cFos could support IAV replication. At the endoplasmic reticulum (ER), cFos activates phospholipid synthesis by physically interacting with the CDP diacylglycerol synthase 1 (CDS1) and phosphatidylinositol 4 kinase type IIα (PI4KIIα) enzymes of the polyphosphoinositide (PIP) lipid pathway [15]. On the contrary, another enzyme involved in the PIP pathway, the CDP diaglycerol inositol 3 phosphatidyltransferase (CDIPT) enzyme, is not regulated by cFos [15]. No major differences in IAV replication were detected upon siRNA-based knockdown of these three enzymes compared to NT-siRNA treated A549 cells (Fig 2A), with cell viability and knockdown efficiency remaining above 95% (S2 Fig). In contrast, upon treatment with T-5224, a specific inhibitor of cFos/AP-1 DNA binding [28], IAV replication was impaired. The viral titers for pH1N1 and WSN viruses were significantly lowered by 0.5 and 2 log_10_, respectively, in A549 cells treated with 20 µM T-5224 compared with DMSO-treated cells (Fig 2B), with cell viability remaining superior to 50% (S3 Fig A). At 6 hpi, cFos was mainly localized in the nucleus, more specifically in the nucleoplasm but not in the nucleoli (marked by anti-fibrillarin), and hardly in the ER (marked by anti-calreticulin) (Fig 2C, S4 Fig).

**Fig 2.**
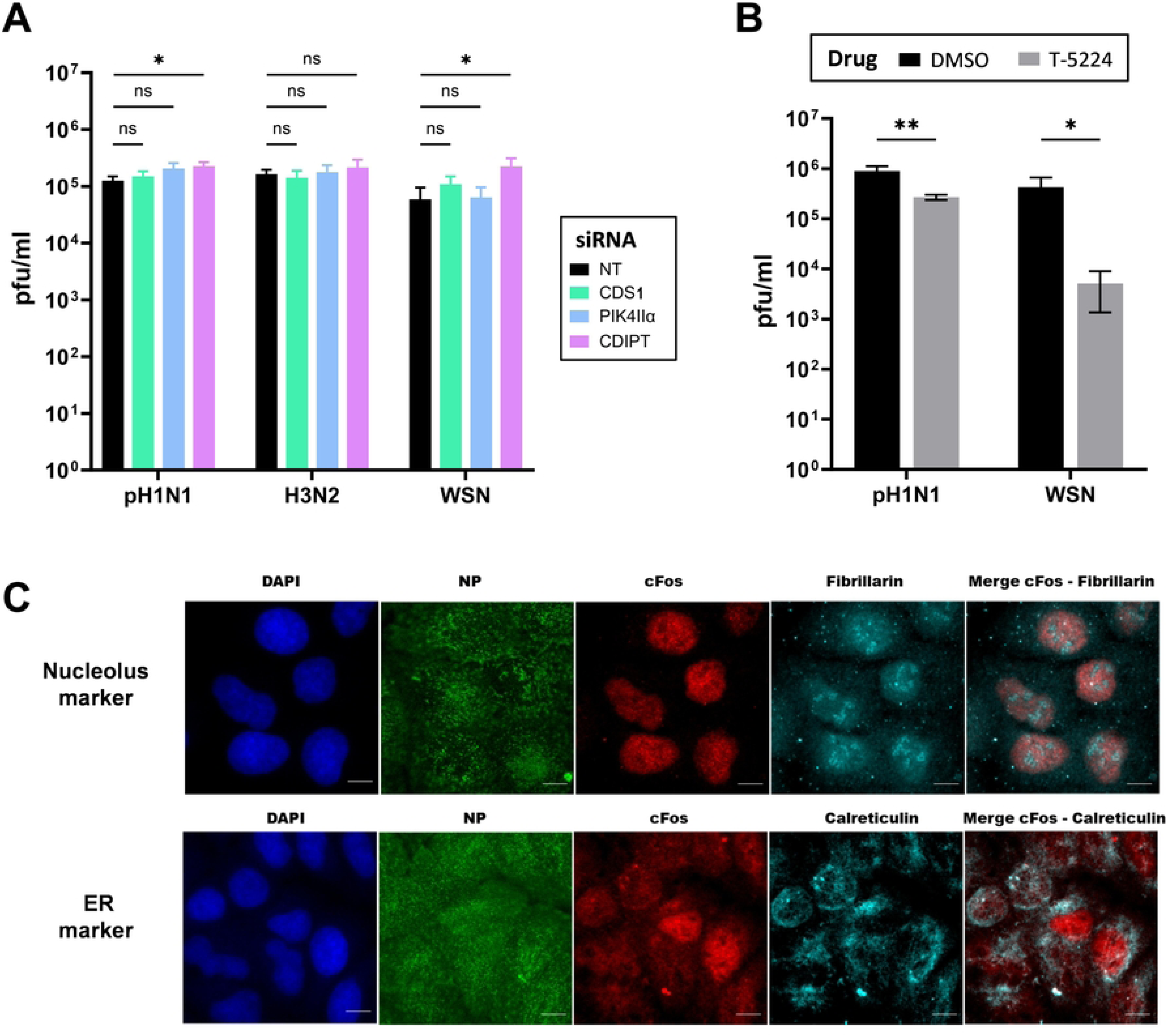
The nuclear function of cFos, but not its cytoplasmic function as an activator of lipid synthesis, appears to regulate IAV multiplication. **(A)** A549 cells were transfected with 25 nM of the non-target (NT) or the indicated siRNAs. At 48 hpt, cells were infected with the following viruses at the indicated moi in pfu/cell: pH1N1, moi of 10^−2^; H3N2, moi of 10^−1^; WSN, moi of 10^−4^. At 24 hpi, viral titers were determined by plaque-forming assay. Results are expressed as the mean ± SD of three independent experiments, and the significance was tested with multiple unpaired t-tests using GraphPad Prism software (ns: not significant, *p< 0.05). **(B)** A549 cells were treated with DMSO (black bars) or 20 µM T5224 (grey bars). At 3h post-treatment, the cells were infected with the following viruses at the indicated moi in pfu/cell: pH1N1, moi of 10^−2^ and WSN, moi of 10^−4^ in presence of DMSO or T5224 20 µM. At 24 hpi, viral titers were determined by plaque-forming assay. Results are expressed as the mean ± SD of three independent experiments, and the significance of the difference to NT was tested with multiple unpaired t-tests using GraphPad Prism software for each virus (*p< 0.05, **p< 0.01). **(C)** Immunofluorescence staining of cFos, fibrillarin (nucleolus marker) and calreticulin (ER marker) in infected A549 cells (WSN, moi of 3). Cells were fixed at 6 hpi, stained with DAPI, and immunostained with anti-NP (infection control), anti-cFos, and anti-fibrillarin or anti-calreticulin. Scale bar = 10 µm.

### The apoptosis is increased in cFos knockdown cells during IAV infection

cFos regulates apoptosis via its nuclear activity. In non-infected cells, upon cFos knockdown or in NT control cells, apoptosis and necrosis rates were about 2% (Fig 3B). Upon IAV infection, both apoptosis and necrosis increased (Fig 3A-B). The apoptosis rate was significantly higher in cFos knockdown cells (12%) compared to control cells (7%). A similar trend was also observed with necrosis, but the difference was not statistically significant (Fig 3A-B).

**Fig 3.**
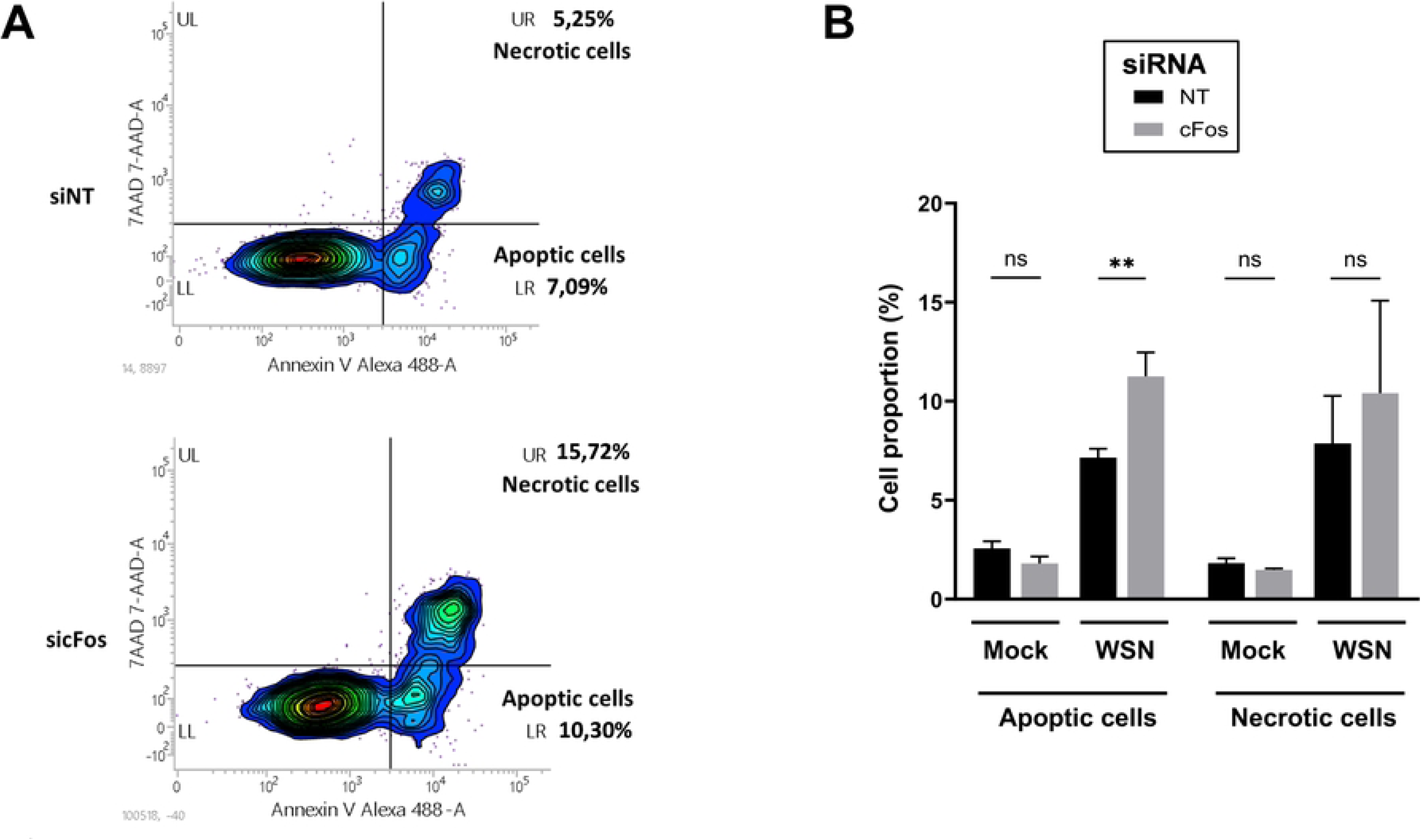
Apoptosis is increased in cFos knockdown cells during IAV infection. A549 cells were treated with 25 nM of the non-target (NT) or cFos siRNAs and infected 48 hpt with WSN at moi of 3 pfu/cell. At 24 hpi, cells were stained with 7AAD (necrotic cells) and Annexin V-AlexaFLuor 488 (apoptotic cells) and subjected to flow cytometry analysis (around 2000 cells analyzed). Mock cells were included in the experiment. **(A)** Contour plot of infected cells treated with NT or cFos siRNA, representative of three independent experiments. UR: necrotic cells. LR: apoptotic cells. **(B)** Relative quantification was performed for necrotic and apoptotic cells. Results are expressed as the mean ± SD of three independent experiments, and the significance of the difference to NT was tested with multiple unpaired t tests using GraphPad Prism software (ns: not significant, **p< 0.01).

### cFos regulates inflammation during IAV infection

To monitor the regulation of cytokine expression by cFos, the expression of inflammatory cytokines IL-1β, IL-6, IL-12, TNF-α (Fig 4A), and type I interferons IFN-α1 and IFN-β (Fig 4B) was measured in cFos knockdown cells during single-cycle IAV infection. IL-1β mRNA induction was significantly lower in cFos knockdown cells compared to the NT-siRNA treated cells (Fig 4A). A decrease in the IL-6 mRNA induction, especially at 3 hpi, was also observed but the overall difference was not statistically significant (Fig 4A). No differences in IL-12A and IL-12B mRNA (both subunits forming the IL-12) as well as TNF-α mRNA were observed between cFos knockdown and NT-siRNA treated cells (Fig 4A). On the contrary, the mRNA level of IFN-β but not of IFN-α1 was significantly higher in cFos knockdown cells (Fig 4B). These findings suggest that, during IAV infection, cFos may promote the transcription of IL-1β and potentially IL-6, and repress that of IFN-β.

**Fig 4.**
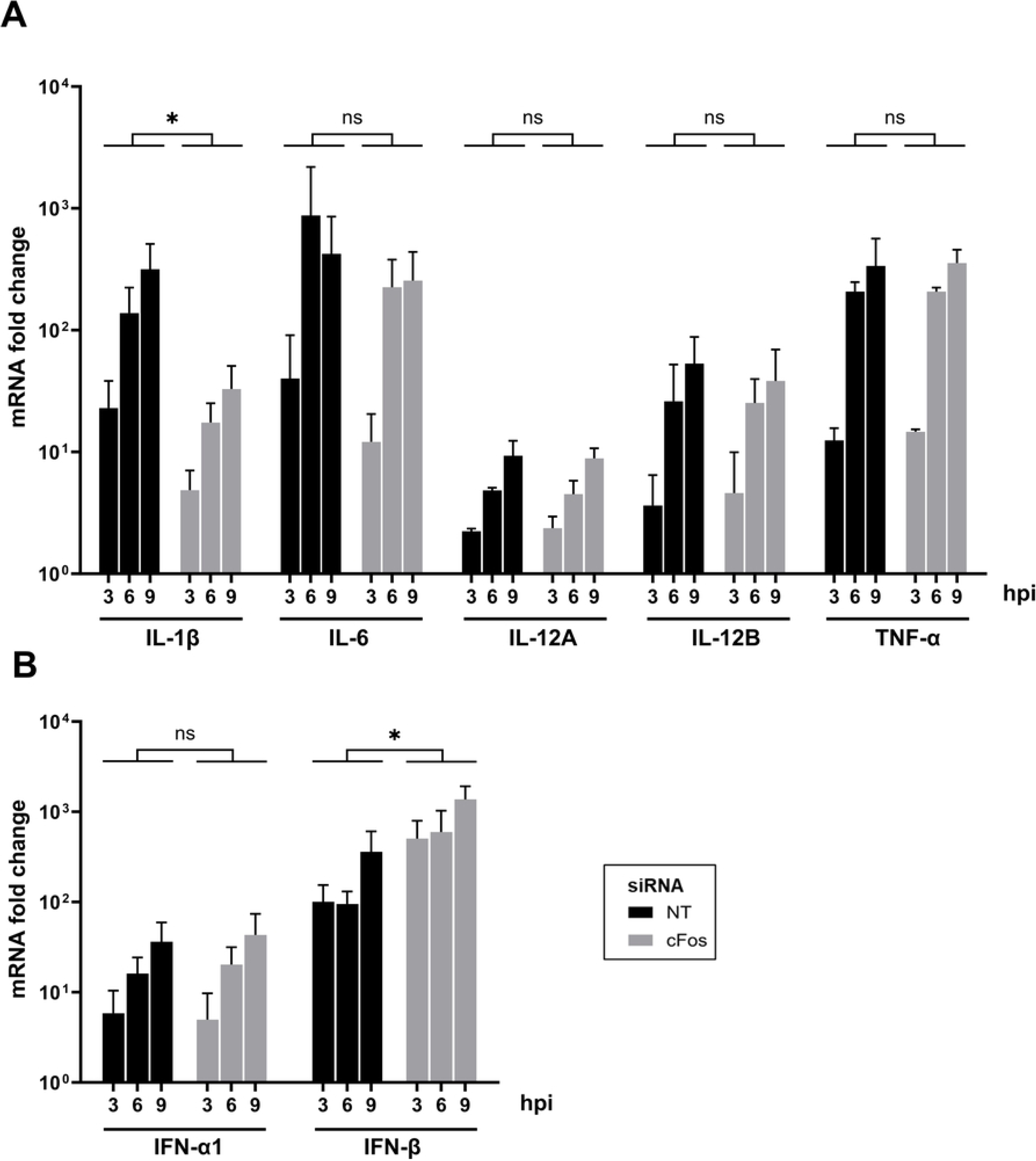
Inflammation and innate immunity are modulated in cFos knockdown cells during IAV infection. A549 cells were treated with 25 nM of the non-target (NT) or cFos siRNAs and infected 48 hpt with WSN at moi of 3 pfu/cell. Total RNAs were extracted at indicated time points and subjected to RT-qPCR specific to **(A)** inflammatory cytokines IL-1β, IL-6, IL12 (IL-12A and IL-12B), and TNF-α, and **(B)** type I interferons IFN-α1 and IFN-β. mRNA fold changes relative to the condition 0 hpi were calculated using the 2^-ΔΔCt^ method. Results are expressed as the mean ± SD of three independent experiments. The area under the curve (AUC) (not shown) was determined for each cytokine and siRNA condition in each independent experiment. The significance of the difference to NT was tested on AUCs with unpaired t tests using GraphPad Prism software (ns: not significant, *p< 0.05).

### Viral transcription and expression of viral proteins are impaired in cFos knockdown cells

The consequences of cFos knockdown on the viral replication were further evaluated. Minigenome assays for three human IAVs - pH1N1, H3N2 and WSN - showed a significant reduction of the viral polymerase activity in cFos knockdown cells (Fig 5A). The levels of vRNA, cRNA, and mRNA production of NP and NA segments during single-cycle WSN infection were further evaluated in both cFos- and NT-siRNA treated cells using strand-specific RT-qPCR [29] (Fig 5B). The production of NA mRNA was lower at 6 hpi (446 vs 2246 RNA fold change, p-value = 0.023) and 9 hpi (162 vs 548 RNA fold change, p-value = 0.104) in cFos knockdown cells compared to NT-siRNA treated cells. On the contrary, the production of vRNA and cRNA of NA tended to be slightly higher at 6 and 9 hpi, although the differences were not significant. For the NP segment, no significant differences were obtained and the levels of mRNA, cRNA and vRNA did not appear to be affected by cFos knockdown (Fig 5B). Similar results were obtained at the protein level. Less NA and M2 viral proteins accumulated at 6 and 9 hpi in cFos knockdown cells than in NT-siRNA treated cells, while no difference in the NP viral protein levels was observed. A slight decrease in NS1 viral protein seemed to occur at 3 hpi upon cFos knockdown but no such differences were observed at later time points. As expected, cFos expression upon IAV infection was increased, especially at 9 hpi in control cells, and knockdown efficiency was confirmed in cFos depleted cells as clearly observed at 6 and 9 hpi (Fig 5C). The same pattern in the expression level of viral proteins was observed upon treatment with T-5224, the specific inhibitor of the cFos/AP-1 dimer DNA binding activity (S3 Fig B). Altogether, these findings highlighted a potential role of cFos in the viral transcription of lately expressed viral proteins.

**Fig 5.**
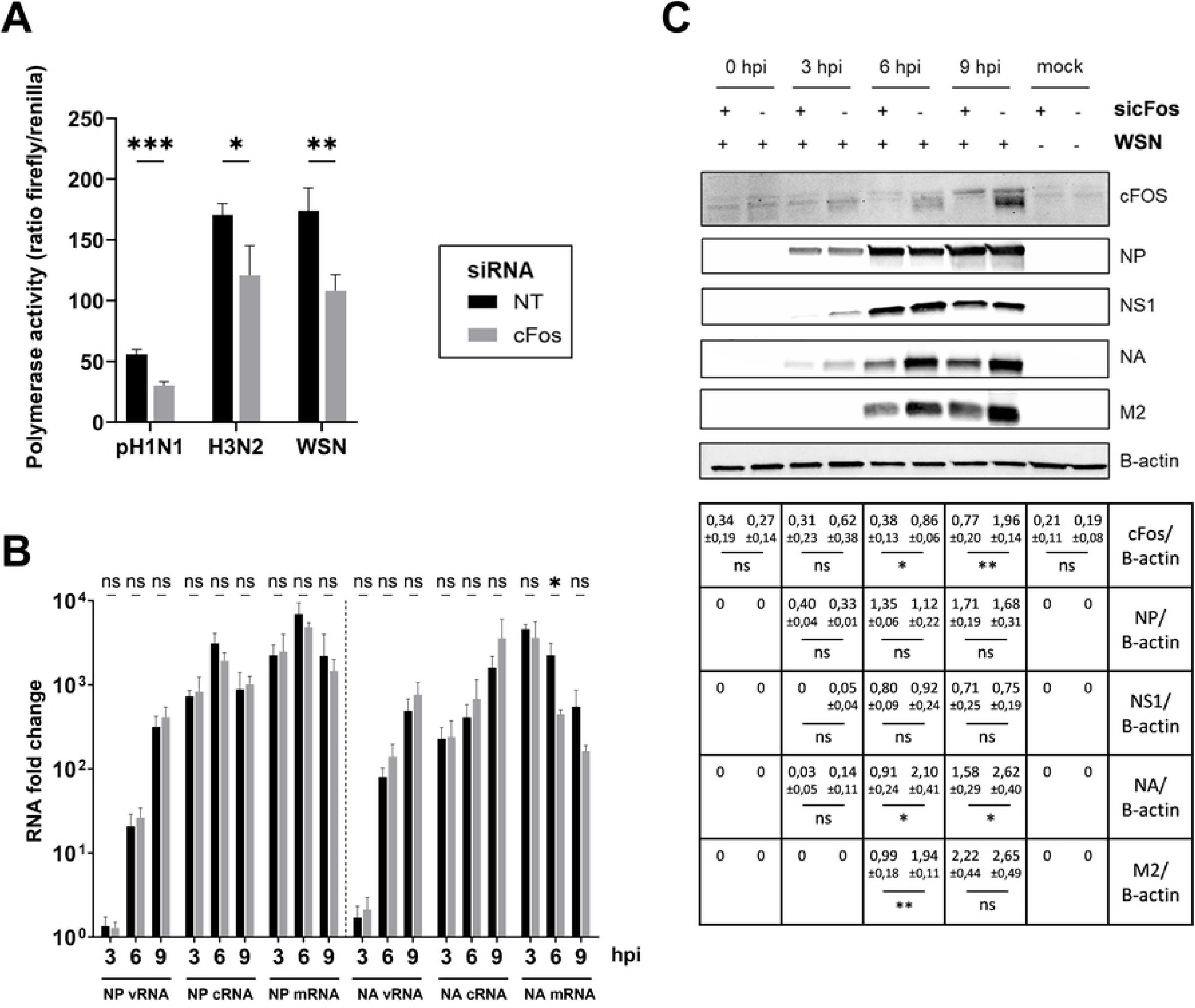
Viral transcription is impaired in cFos knockdown cells. **(A)** HEK293T cells treated during 48h with 25 nM of non-treated NT or cFos siRNAs were transfected with minigenome components. Firefly and Renilla luciferase activities were measured 24 hpt using the Dual-Glo Luciferase assay. Firefly luciferase activity (proportional to viral polymerase activity) was normalized to Renilla luciferase activity to take into account the transfection rate and cell density. Results are expressed as the mean ratio ± SD of three independent experiments. The significance of the difference to NT was tested by an unpaired t-test in GraphPad Prism Software (*p< 0.05, **p< 0.01, ***p< 0.001). **(B,C)** A549 cells were treated with 25 nM of NT or cFos siRNAs and infected 48 hpt with WSN at moi of 3 pfu/cell. **(B)** Total RNAs were extracted at indicated time points and subjected to strand-specific RT-qPCR [29]. RNA fold changes relative to the condition 0 hpi were calculated using the 2^-ΔΔCt^ method. The results are expressed as the mean ± SD of three independent experiments. The significance of the difference to NT was tested with unpaired t tests using GraphPad Prism software (ns: non significant, *p< 0.05). **(C)** Total cell lysates were harvested at the indicated times post-infection and analyzed by immunoblot using antibodies directed against the indicated proteins. Immunoblot results representative of three independent experiments are shown. Band intensity of the indicated proteins was normalized to β-actin, and the mean ratios ± SD of three independent experiments are presented in the table. The significance of the difference to NT (indicated by ‘-‘ in sicFos) was tested by unpaired t-tests in GraphPad Prism Software (ns: non significant, *p< 0.05, **p< 0.01).

## Discussion

AP-1 transcription factors are commonly induced by viral infections and were found to regulate the replication of several viruses [16–19]. IAV infection was shown to activate AP-1 transcription factors through phosphorylation by the JNK signaling pathway [24]. However, the role of AP-1 in IAV replication remains unclear. In the current study, we found that the transcription of cFos and cJun AP-1 factors was upregulated following IAV infection and that cFos appeared to support IAV replication.

The role of cFos is dependent on its location in the cell. At the endoplasmic reticulum, cFos activates *de novo* phosphatidylinositol phosphate (PIP) lipid synthesis by interacting with CDS1 and PI4KIIα enzymes [15]. The individual siRNA-based knockdown of CDS1 and PI4KIIα enzymes did not impair IAV replication (Fig 2A). Although the individual knockdown of CDS1 in cardiomyoblast cells was previously shown to be critical and to impair the PIP pathway [30], it cannot be ruled out that CDS1 and PI4KIIα protein isoforms might have played a compensatory role in our experiments. However, it is more likely that cFos does not influence IAV replication through its cytoplasmic role as an activator of PIP lipid synthesis, but rather through its nuclear role as an AP-1 transcription factor. This is further supported by the localization of cFos in the nucleoplasm during IAV infection (Fig 2C – S4 Fig) and the reduction of IAV replication in the presence of cFos nuclear activity inhibitor T-5224 (Fig 2B). The most described cFos AP-1 dimer nuclear partner, cJun, did not appear to regulate IAV replication in our study, although it was previously found to support H5N1 IAV replication in A549 cells [23]. These discrepancies may be attributed to differences in the viral strains or to the presence of residual cJun, since in our study, cJun depletion was less effective compared to cFos depletion. cFos might also regulate viral replication independently of its partner cJun, through binding to nucleic acids. In this regard, the overexpression of cFos mRNA upon infection may increase the likelihood of cFos forming homodimers or remaining as monomers, which were both shown to be able to bind cellular DNA [31,32]. The activity of cFos is regulated via phosphorylation by MAPK kinases, including ERKs and p38 [33,34]. These kinases are activated during IAV infection [35,36]. ERK5 was shown to phosphorylate cFos at the serine 32 position to increase its stability and nuclear localization [37]. Unfortunately, our experimental setup did not allow us to distinguish between the phosphorylated (Ser-32, using a specific antibody) and total cFos, and therefore to determine whether cFos phosphorylation was necessary for the observed effect on the virus replication.

The role of cFos on apoptosis is unclear, with both pro- and anti-apoptotic roles mainly observed in cancer models [7,9,38]. IAV was shown to protect cells from premature apoptosis [39]. However, at the late stage of infection, IAV promotes apoptosis to facilitate the spread of viral particles to neighboring cells [39]. In our study, cFos knockdown significantly increased apoptosis in IAV-infected A549 cells (Fig 3). A similar increase of apoptosis was also observed in cFos knockdown H1299 human epithelial lung cells infected with a gamma coronavirus [19]. Therefore, during viral infection, cFos seems to be involved in cell survival by inhibiting apoptosis.

Although certain transcription factors with known antiviral functions, such as STAT-1 and NF-Kb p65, have been shown to facilitate IAV replication [40,41], the underlying molecular mechanisms remain poorly understood. AP-1 factors are widely recognized as transcriptional activators that promote expression of pro-inflammatory cytokines (reviewed in ref. [2]) and interferon-β and -γ [42,43]. Our results suggest that cFos may counteract cellular antiviral response by downregulating virus-induced IFN-β expression (Fig 4B). Activation of the *IFNb1* gene transcription occurs through a signaling cascade that requires the cooperative binding of ATF2/cJun AP-1 dimer [42]. Since cFos possesses a higher affinity for cJun than ATF2 and was shown to displace the ATF2/cJun dimer [44], cFos overexpression upon IAV infection could decrease cJun availability for ATF2, leading to a reduction in IFN-β transcription. The absence of the IFN-β expression or the inhibition of IFN-β activation by the cellular exonuclease XRN1 was shown to facilitate IAV replication [45,46]. IFN-β triggers the transcriptional activation of antiviral interferon-stimulated genes (ISGs). Given that these ISGs were shown to be upregulated at early stage (4 hpi) during IAV infection in epithelial cells [47], enhanced ISG expression resulting from increased IFN-β levels could potentially account for the impaired viral transcription of NA mRNA observed in cFos knockdown cells during single-cycle IAV infection experiments (Fig 5).

The impaired viral transcription of NA mRNA (Fig 5) may also suggest a direct role of cFos in the viral transcription. IAV relies on cap-snatching to obtain host-capped RNA fragments to initiate its own transcription, a process mediated by the interaction between the viral polymerase complex and the cellular DNA-dependent RNA polymerase II (RNAP II) [48]. cFos/cJun AP-1 dimer was shown to bind enhancer sequences, and then to recruit the chromatin remodeling BRG1-Associated Factor (BAF) complex, leading to an accessible chromatin state [49]. Interaction between the Brg1 subunit of the BAF complex and the RNAP II was further shown to participate in the formation of the transcriptional pre-initiation complex [50]. As a transcription factor, cFos may thus facilitate the accessibility to the RNAP II for the viral polymerase complex, therefore helping in the cap-snatching process. Another chromatin remodeling complex was shown to support IAV viral transcription through direct interaction with the viral polymerase complex [51]. Using a split luciferase protein complementation assay, no direct interaction between cFos and the viral polymerase complex could be shown (S5 Fig), indicating that cFos would act through interaction with one or several additional cellular partners, or with a viral protein other than PB2, PB1 or PA polymerase subunits. Both IAV viral transcription and replication take place in the nucleus, and a conformational change of the viral polymerase is required to switch from transcription to replication status [52]. Interestingly, in parallel to the decrease of NA mRNA levels in cFos knockdown cells, an increase of NA vRNA and cRNA levels seemed to occur, suggesting that cFos could be involved in the regulation of the timing of the viral transcription/replication switch.

In conclusion, this study highlighted cFos as a pro-viral factor that facilitates IAV replication. cFos was observed to extend cell survival and downregulate IFN-β expression during IAV infection, potentially promoting viral replication. In addition, cFos may be mechanistically involved in viral transcription. However, the exact mechanisms by which cFos support IAV replication remain unclear. It also remains enigmatic why the decrease observed in the transcription and expression of viral proteins only affected some viral proteins, especially those that are lately expressed in the virus cycle.

## Materials and Methods

### Cell lines

HEK293T, A549, MDCK, and MDCK-SIAT1 [53] cells were provided by Institut Pasteur Paris. HEK293T and A549 cells were grown in Dulbecco’s modified Eagle’s medium (DMEM) (Life Technologies cat#41965039) supplemented with 10% fetal bovine serum (FBS) (Life Technologies cat#10270-106), 10 U/ml of penicillin and 10 µg/ml of streptomycin (Life Technologies cat#15140122). MDCK and MDCK-SIAT1 cells were grown in Modified Eagle’s Medium (MEM) (Life Technologies cat#31095029) supplemented with 5% FBS, 10 U/ml of penicillin and 10 µg/ml of streptomycin. All cell lines were grown at 37°C in 5% CO_2_.

### Viruses

Three human IAVs were used: the two currently circulating seasonal IAVs in human population, A/Bretagne/7608/2009 (H1N1)pdm09 (referred to as pH1N1 in Results) and A/Centre/1003/2012(H3N2); and the laboratory-adapted A/WSN/33 (H1N1) (referred to as WSN in Results). All viruses were produced by reverse genetics [54,55] and amplified on MDCK cells, except for the H3N2 strain, which was amplified on MDCK-SIAT1 cells. pH1N1 and H3N2 subtypes grow poorly in A549 cells. Two mutations in the pH1N1 HA (A517G, G834A) and one mutation in the H3N2 HA (G460T) were introduced, by site-directed mutagenesis, into the HA-encoding plasmid pH1N1 and H3N2 of the reverse genetic system to generate A549-adapted pH1N1 and H3N2 strains, capable of efficient replication in A549 cells [56].

### Antibodies, chemicals, and reagents

Rabbit anti-β-actin (cat#4967), rabbit anti-cFos (cat#2250), goat anti-rabbit IgG, HRP-linked (cat#7074), and horse anti-mouse IgG, HRP-linked (cat#7076) antibodies were purchased from Cell Signaling Technology. Mouse anti-M2 (cat#MA1-082), rabbit anti-NP (cat#PA5-32242), chicken anti-calreticulin (cat#PA1-902A), chicken anti-fibrillarin (cat#PA5-143565A), goat anti-rabbit Alexa Fluor Plus 647 (cat#A32733), and goat anti-chicken IgY (H+L) AlexaFluor 594 (cat#A-11042) antibodies were purchased from ThermoFisher Scientific. The rabbit anti-NA antibody (cat#GTX125974) was purchased from Genetex. The mouse anti-NS1 (cat#sc-130568) and the mouse anti-NP (cat#sc-80481) antibodies were purchased from Santa Cruz Biotechnology. The goat anti-mouse IgG-FITC antibody (cat#F0257) was purchased from Sigma-Aldrich. The Annexin V-Alexa Fluor 488 apoptosis detection reagent (cat#A13201) was purchased from ThermoFisher Scientific, and the 7-Amino-Actinomycin D (7-AAD) (cat#559925) was purchased from BD Biosciences. The T-5224 small inhibitor (cat#HY-12270) was purchased from MedChemExpress. CellTiter-Glo® Luminescent Cell Viability Assay (cat#G7570), Dual-Glo® Luciferase Assay System (cat#E2920), and Renilla Luciferase Assay System (cat#E2810) were purchased from Promega.

### siRNA-based assays

#### Multi-cycle infection

siRNAs were purchased from Horizon Discovery (ON-TARGETplus SMARTpools and Non-targeting Control pool). The siRNA sequences were designed by Horizon Discovery and are listed in S1 table. 6.5 µl of siRNA at a concentration of 5 µM was added to 243.5 µl of DharmaFECT1 transfection reagent (Horizon discovery cat#T-2001) and OptiMEM GlutaMAX medium (Life Technologies cat#51985034) to obtain a final volume of 250 µl. The 250 µl mixture was added to one well of a 24-well tissue culture plate (Greiner). Following a 30-minute incubation period at room temperature, 2×10^5^ A549 cells diluted in 1 ml of DMEM supplemented with 5% FBS were seeded on the top of siRNA-transfection reagent complexes and incubated at 37°C in 5% CO_2._ The final concentration of siRNA was 25 nM per well. At 48 hours post transfection (hpt), the cells were washed and infected with pH1N1 at multiplicity of infection (moi) of 10^−2^ or 10^-3^, WSN at moi of 10^−4^, and H3N2 at moi of 10^−1^ or 10^−2^ at 35°C (5% CO_2_). Supernatants were collected at 0, 24 and/or 48 hours post infection (hpi), and viral titers were determined by plaque assays using MDCK-SIAT1 cells [57].

#### Single-cycle infection

The cells were treated with siRNAs as described above. At 48 hpt, the cells were washed, inoculated with WSN at moi of 3, and incubated for 1 h at room temperature. Inoculum was then removed, fresh OptiMEM GlutaMAX medium was added, and cells were incubated at 35°C (5% CO_2_). Timepoint 0 hpi was defined as the start of the cell incubation at 35°C.

#### Cell viability and luciferase-based knockdown efficiency experiments

siRNA reverse transfection was performed in a white 96-well tissue culture plate (Greiner). Volumes were adjusted to obtain a siRNA final concentration of 25 nM per well. Briefly, 0.65 µl of siRNA at a concentration of 5 µM was added to 24.35 µl of DharmaFECT1 transfection reagent and OptiMEM GlutaMAX medium to obtain a final volume of 25 µl. The 25 µl mixture of siRNA-DharmaFECT1 was added to one well of the white 96-well plate. Following a 30-minute incubation period at room temperature, 1.5×10^4^ A549 cells diluted in 100 µl of DMEM supplemented with 5% FBS were seeded on top of the siRNA-transfection reagent complexes and incubated at 37°C in 5% CO_2_. Cell viability was determined at 48 hpt using the CellTiter-Glo Luminescent Viability Assay kit according to the manufacturer’s instructions (Promega). To measure the efficiency of siRNA-knockdown, siRNA-treated A549 cells were transfected at 24 h post-siRNA transfection with 10 ng of plasmids encoding siRNA-targeted protein fused with the full-length Gaussia luciferase (pGlucFL) using lipofectamine 3000 (ThermoFisher Scientific cat#L3000001). The luciferase activity was measured 24h later in cell lysates using the Renilla luciferase assay reagent (Promega). All luciferase activities were measured on a GloMax Explorer (Promega).

### RT-qPCR

RNAs were extracted using the Quick-RNA Miniprep Kit (Zymo Research cat#R1054) from A549 cells treated with siRNA and infected with WSN or mock infected. The kit uses a column to remove most of the genomic DNA and a subsequent treatment with DNase I to degrade the remaining genomic DNA. RNA concentrations were measured with Nanodrop One (Thermo Fischer Scientific) and adjusted to 100 ng/µL. 200 ng RNA was reverse transcribed using the Accuscript High Fidelity 1st strand cDNA synthesis kit (Agilent cat#200820) following the manufacturer’s instructions. Briefly, 200 µg RNA was mixed with 4 µl buffer 10x AccuScript High Fidelity RT-PCR System, 1,6 µl 100 mM dNTPs, 1 µg anchored-oligo(dT) primer, and nuclease-free water in a 33 µL reaction mixture. The mixture was heated for 5 min at 65°C and cooled for 5 min to room temperature before the addition of 40 U RNase block, 10 mM DTT, and 2 µl Accuscript RT in a 40 µl final volume reaction mixture. Reverse transcription was performed at 42°C for 60 min in a thermocycler, and the RT enzyme was then inactivated at 72°C for 15 min. Real-time qPCR (qPCR) was performed using GoTaq qPCR master mix containing BRYT green dye (Promega cat#A6001) on an ARIA MX (Stratagene MX3005P, Agilent technologies) in a total volume of 20 µl. Each reaction mixture included 10 µL GoTaq qPCR MasterMix, 1 µL of each diluted forward and reverse primers (10 µM), 1.25 µL template cDNA, and nuclease-free water up to 20 µL. Primers for the AP-1 transcription factors (sequences based on ref. [19]) were purchased from Integrated DNA Technologies. Primer sequences for cytokine genes and phospholipid enzymes were designed and verified by Sigma-Aldrich. All the primers are listed in S2 Table. A control with 1.25 µL of water instead of cDNA was included in each run to check for reagent contamination. A “No RT” control was also included for each targeted gene of the qRT-PCR experiments to check the efficiency of the DNase I treatment. The thermal cycling conditions for all qPCRs were as follows: denaturation at 95°C for 2 min, followed by 40 cycles consisting in 95°C for 15 sec, and an annealing and extension phase at 60°C for 1 min. Afterwards, a melting curve analysis was performed to determine the specificity of the reaction products. The modulation of RNA expression for all targeted genes was normalized to β-actin expression (housekeeping gene) and analyzed using the 2^-ΔΔCt^ method (delta-delta Ct method).

### Strand-specific RT-qPCR

RNAs were extracted using the Quick-RNA Miniprep Kit (Zymo Research cat#R1054) from A549 cells treated with siRNA and infected with WSN or mock infected. Strand-specific RT-qPCR for NP and NA vRNAs, cRNAs, and mRNAs was performed as previously described [29]. Briefly, cDNAs complementary to NP and NA vRNAs, cRNAs, and mRNAs were synthesized using SuperScript™ IV Reverse Transcriptase (ThermoFisher Scientific cat#18090010) and tagged primers in order to add a strand-specific tag unrelated to influenza virus sequence at the 5′ end. Tagged cDNAs were then used as a template for the qPCR reaction using a tag-specific primer and a segment-specific primer. β-actin (housekeeping gene) mRNA was reverse transcribed using anchored-oligo(dT) and SuperScript™ IV Reverse Transcriptase (ThermoFisher Scientific cat#18090010). All qPCRs were performed using GoTaq qPCR master mix containing BRYT green dye (Promega cat#A6001) on an ARIA MX (Stratagene MX3005P, Agilent technologies) in a total volume of 20 µl. Each reaction mixture included 10 µL GoTaq qPCR MasterMix, 1 µL of each diluted forward and reverse primers (10 µM), 7 µL 10-fold-diluted template cDNA, and nuclease-free water up to 20 µL. The primers were designed as described in ref. [29] to specifically detect NP and NA vRNAs, cRNAs and mRNAs. A control with 7 µL of water instead of cDNA was included in each run to check for reagent contamination. The thermal cycling conditions for all qPCRs were as follows: denaturation at 95°C for 2 min, followed by 50 cycles consisting in 95°C for 15 sec, and an annealing and extension phase at 60°C for 1 min. Afterwards, a melting curve analysis was performed to determine the specificity of the reaction products. The modulation of vRNA, cRNA and mRNA expression for all targeted genes was normalized to β-actin expression (housekeeping gene), and RNA fold changes relative to the condition 0 hpi were calculated using the 2^-ΔΔCt^ method. All the primers for both reverse transcription and qPCR are listed in S3 Table.

### Western blot

A549 cells treated with siRNA and infected with WSN or mock infected were lysed at 0, 3, 6, and 9 hpi using RIPA lysis and extraction buffer (ThermoFisher Scientific cat#89900) supplemented with 100-fold diluted Halt™ Protease and Phosphatase Inhibitor Cocktail, EDTA-free (ThermoFisher Scientific cat#78441). Briefly, the cells were washed with cold PBS, and 200 µl supplemented RIPA buffer was added to each well of the 12-well plate. Following a 15 min incubation period on ice, lysed cells were collected and centrifuged at 14,000 g for 15 min. Cell lysate supernatants were kept, and protein quantification was performed using Pierce™ BCA Protein Assay Kits (ThermoFisher Scientific cat#23225) following manufacturer’s instructions. Cell lysates were then mixed with 4x Laemmli Sample Buffer (BioRad cat#1610747) supplemented with 355 mM of β-mercaptoethanol (Sigma-Aldrich cat#M6250) and boiled at 90°C for 10 min. Equal amounts of protein samples were loaded into each well and separated using 8–16% Mini-PROTEAN® TGX Stain-Free™ Protein Gel (BioRad cat#4568105) in a Mini-PROTEAN Tetra Vertical Electrophoresis Cell (BioRad cat#1658004), at 200 V for 30 min. The resolved proteins were then transferred to a 0.2 µM nitrocellulose membrane for 7 min at 25 V using the Trans-Blot® Turbo™ Transfer System (BioRad cat#1704150). The membrane was blocked for 1 h at room temperature with PBS-0.05% Tween 20-3% BSA, then incubated with indicated primary antibodies diluted in PBS-0.05% Tween 20-3% BSA at 4°C overnight. After 3 washes for 5 minutes with PBS-0.1% Tween 20, the membrane was incubated with 1:5000 HRP-linked goat anti-rabbit IgG or HRP-linked horse anti-mouse IgG, at room temperature for 2 hours. After 3 washes for 5 minutes with PBS-0.1% Tween 20, proteins were detected by chemiluminescence using Pierce™ ECL Plus Western Blotting Substrate (ThermoFisher Scientific cat#32132) and ChemiDoc imaging system (Biorad). Protein band intensities were determined using ImageJ software. All experiments were repeated three times, with similar results, and one of the representative immunoblots is shown.

### Immunofluorescence staining

A549 (2×10^5^) cells were seeded on coverslips (13 mm of diameter) in 24-well plate (Greiner) in DMEM supplemented with 10% FBS, 10 U/ml of penicillin and 10 µg/ml streptomycin, and incubated at 37°C for 24 h. Cells were infected with WSN at moi of 3 pfu/cell. At 0, 3, 6, and 9 hpi, cells were fixed with 3% paraformaldehyde for 20 min and permeabilized with PBS-0.1% Triton X100 for 20 min. Cells were blocked with PBS-3% BSA for 1 hour and incubated with primary antibodies anti-cFos (1/200), anti-NP (1/200), and anti-fibrillarin (1/200) or anti-calreticulin (1/100) overnight at 4°C. Following multiple washes with PBS, cells were incubated with FITC goat anti-mouse IgG (1/200), Alexa Fluor 594 goat anti-chicken IgY (1/200), and Alexa Fluor 647 goat anti-rabbit (1/300) secondary antibodies for 1h at room temperature. Following multiple washes with PBS, coverslips were mounted in ProLong Gold Antifade Mountant with DNA Stain DAPI (ThermoFisher Scientific cat#P36941) and analyzed under an inverted fluorescence microscope (Nikon ECLIPSE Ts2R) using an x60 objective lens.

### Minigenome assay

HEK293T (3×10^4^) cells were transfected with NT or cFos siRNA in a white 96-well tissue culture plate, as described above. After 48h of knockdown, cells were transfected using Polyethyleneimine « MAX » (MW 40,000) 1 mg/ml (Polysciences cat#24765) with expression pCIneo or pcDNA3 plasmid vectors encoding the IAV proteins PA, PB1, PB2 (25 ng each plasmid) and NP (50 ng) from different IAV strains (pH1N1, WSN, H3N2), a reporter plasmid vector (pPR7-firefly-(-)) encoding the firefly luciferase in the negative-sense orientation flanked by the noncoding regions of the segment 5 of WSN driven by a polymerase I [Pol I] promoter (5 ng) [58], and a plasmid vector (polIII-Renilla) constitutively expressing Renilla luciferase (5 ng). Twenty-four hours post-transfection, luciferase activities were measured using the Dual-Glo luciferase assay system (Promega). Polymerase activity, proportional to Firefly luciferase activity, was normalized to Renilla luciferase activity to take into account the transfection rate.

### Measure of apoptosis and necrosis rates

A549 cells were transfected with NT or cFos siRNA as described above. After 48h of knockdown, cells were infected with WSN at moi of 3 pfu/cell or mock infected. At 24 hpi, the apoptosis and necrosis rates were determined. Cells were washed with PBS and detached using trypsin-EDTA 0.05% phenol red (ThermoFisher Scientific cat#25300054). DMEM supplemented with 10% FBS was then added, and the collected cells were centrifuged at 2000 rpm for 5 min. Pellet cells were resuspended in Annexin-binding buffer (HEPES 10mM, NaCl 140 mM, CaCl_2_ 2.5 mM, pH 7.4) at 10^6^ cells/ml. 100 µl of cells was then transferred into a FACS tube, and 5 µl of Annexin V-Alexa Fluor 488 apoptosis detection reagent and 7-AAD was added. Staining was performed for 15 min at room temperature. 400 µl of Annexin-binding buffer was then added. Cells were further fixed in 4% paraformaldehyde at 4°C in the dark until analysis. As a positive control, cells were also treated with 10 µM Camptothecin (ThermoFisher Scientific cat#J62523.MD), known to induce apoptosis. Cells were acquired on a FACSVerse flow cytometer (BD Biosciences). Data was analyzed with the FACSuite software (BD Biosciences). Around 2000 events were counted for each sample.

### Statistical analysis

Unpaired Student’s t-test was used for all statistical analyses, using GraphPad Prism software v.9.00 (GraphPad Software). Differences between groups were considered statistically significant at p< 0.05; significance levels are as follow: *p< 0.05, **p< 0.01, ***p< 0.001, ns: non significant.

## Acknowledgments

We thank Samuel Kindylides and Brecht Droesbeke for support with the microscopy experiments, and Celine De Sterck for technical assistance with the immunoblots.

## Supporting information

**S1 Fig. Cell viability (A) and knockdown efficiency (B) upon siRNA transfection. (C) Control RAB11A siRNA treatment reduced IAV replication.** A549 cells were transfected with 25 nM of the NT or the indicated siRNAs. **(A)** Cell viability was determined at 48 hpt using the CellTiter-Glo Luminescent Viability Assay. Luciferase activities in the indicated siRNAs were compared to the non-target (NT) treated cells to determine a percentage of viability. siPLK1 was included as a known cytotoxic control. The results are expressed as the mean percentages ± SD of three independent experiments. The significance of the difference to NT was tested with unpaired t tests using GraphPad Prism software (ns: non significant, **p< 0.01). **(B)** At 24h post siRNA transfection, a second transfection with plasmids encoding cFos, cJun and RAB11A proteins fused with the full-length Gaussia luciferase (pGlucFL) was performed to assess knockdown efficiency. Ratios of the luciferase activities obtained in cells transfected with the indicated siRNAs to the ones obtained in cells transfected with the NT siRNA are shown. The results are expressed as the mean percentages ± SD of three independent experiments. The dashed line corresponds to a relative protein expression of 50% in silenced cells compared to NT-treated cells. **(C)** At 48 hpt, cells were infected with the following viruses at the indicated moi in pfu/cell: pH1N1, moi of 10^−3^; H3N2, moi of 10^−2^; WSN, moi of 10^−4^. At 24 hpi, viral titers were determined by plaque-forming assay. Results are expressed as the mean ± SD pfu/ml of three independent experiments. The significance of the difference to NT was tested with unpaired t tests using GraphPad Prism software (ns: non significant, *p< 0.05, ***p< 0.001).

**S2 Fig. Cell viability and knockdown efficiency upon siRNA knockdown of lipid synthesis enzymes.** A549 cells were transfected with 25 nM of the indicated siRNAs **(A)** Cell viability was determined at 48 hpt using the CellTiter-Glo Luminescent Viability Assay. Luciferase activities in the indicated siRNAs and the non-target (NT) treated cells were compared to determine a percentage of viability. Results are expressed as the mean percentages ± SD of three independent experiments. **(B)** Total RNAs were extracted at 48 hpt and the mRNA expression level of each gene was determined by RT-qPCR. Results are expressed at the mean ± SD of the percentage of remaining mRNA in each siRNA condition, as compared to NT control (calculated using the 2^-ΔΔCt^ method), determined in three independent experiments.

**S3 Fig. Inhibition of AP-1 transcriptional activity of c-Fos/cJun AP-1 by the T-5224 inhibitor impaired viral protein expression.** A549 cells were treated with DMSO or 20 µM T5224. **(A)** Cell viability was determined at 24 h post-treatment using the CellTiter-Glo Luminescent Viability Assay. Luciferase activities in the T-5224 condition were compared to the DMSO condition to determine a percentage of viability. The results are expressed as the mean percentages ± SD of three independent experiments. **(B)** At 16 h post-treatment, cells were infected with WSN at moi of 3 pfu/cell in presence of T-5224 20µM or DMSO. Total cell lysates were harvested at the indicated times post-infection and analyzed by immunoblot using antibodies directed against the indicated proteins. Band intensity of the indicated proteins was normalized to ß-actin and the mean ratios ± SD of three independent experiments are presented in the table. The significance of the difference to DMSO (indicated as ‘-‘ in T-5224) was tested by an unpaired t-test in GraphPad Prism Software (ns: non significant, *p< 0.05, **p< 0.01, ***p<0.001).

**S4 Fig. Immunofluorescence staining of cFos in A549 cells during single cycle infection (WSN, moi of 3).** Cells were fixed at 0, 3, 6 and 9 hpi, stained with DAPI and immuno-stained with anti-NP (infection control), and anti-cFos antibodies. Scale bar = 10 µm.

**S5 Fig. Split luciferase protein complementation assay between the IAV viral polymerase complex and cFos.** HEK293T cells were transfected with plasmids encoding PA, PB1, PB2 fused to Gluc1 split domain of the Gaussia Luciferase, and cellular proteins (cFos, hANP32, ATXN1, TBP, or GTF2B) fused to Gluc2 split domain of the Gaussia Luciferase. At 24 hpt, cells were lysed, and luminescence was measured (RLU). Normalized luminescence ratio (NLR) was calculated for each combination as follows: NLR = RLU (A-Gluc1 + B-Gluc 2) / [RLU (A-Gluc1 + Gluc2) + RLU (Gluc1 + B-Gluc2)]. The human ANP32 (hANP32), described to interact with the polymerase complex, is considered as a positive control (green bar) while ATXN1, TBP, and GTF2B are random proteins (negative control - red bars). The results are expressed as the mean percentages ± SD of three independent experiments. The significance of the difference to cFOS was tested by unpaired t-tests in GraphPad Prism Software (ns: non significant, ***p< 0.001).

**S1 Table. siRNA sequences**

**S2 Table. RT-qPCR primer sequences (AP-1 primer sequences from Yuan et al. 2021)**

**S3 Table. Strand-specific RT-qPCR primer sequences (from Kawakami et al. 2011)**

